# Potassium homeostasis during disease progression of Alzheimer’s Disease

**DOI:** 10.1101/2024.05.23.595252

**Authors:** Evgeniia Samokhina, Armaan Mangat, Chandra S. Malladi, Erika Gyengesi, John W. Morley, Yossi Buskila

## Abstract

Alzheimer’s disease (AD) is an age-dependent neurodegenerative disorder characterized by neuronal loss leading to dementia and ultimately death. Whilst the loss of neurons is central to the disease, it is becoming clear that glia, specifically astrocytes, contribute to the onset and progression of neurodegeneration. Astrocytic role in retaining ion homeostasis in the extracellular milieu is fundamental for multiple brain functions, including synaptic plasticity and neuronal excitability, which are compromised during AD and affect neuronal signalling. In this study, we have measured the astrocytic K^+^ clearance rate in the hippocampus and somatosensory cortex of a mouse model for AD during disease progression. Our results establish that astrocytic [K^+^]_o_ clearance in the hippocampus is reduced in symptomatic 5xFAD mice, and this decrease is region-specific. The decrease in the [K^+^]_o_ clearance rate correlated with a significant reduction in the expression and conductivity of Kir4.1 channels and a decline in the number of primary connected astrocytes. Moreover, astrocytes in the hippocampus of symptomatic 5xFAD mice demonstrated increased reactivity which was accompanied by an increased excitability and altered spiking profile of nearby neurons. These findings indicate that the supportive function astrocytes typically provide to nearby neurons is diminished during disease progression, which affects the neuronal circuit signalling in this area and provides a potential explanation for the increased vulnerability of neurons in AD.

## Introduction

Alzheimer’s disease (AD) is an irreversible, progressive neurological disorder characterized by memory loss and impairment of cognitive functions. Described for the first time in 1906 by Alois Alzheimer, the major neuropathological markers found in AD brains are extracellular amyloid-beta (Aβ) plaques and intracellular neurofibrillary tangles composed of hyper and abnormally phosphorylated tau protein (1). These pathologic features are accompanied by gliosis and widespread neuronal loss (2). However, prior to the extensive neurodegeneration, neurons in AD patients and transgenic AD mouse models, express aberrant patterns of cellular and network oscillatory rhythmic activity that may facilitate the cognitive decline associated with AD (3). Some of the suggested underlying mechanisms for the abnormal network activity includes the disruption of synaptic functionality (4) and neuronal hyperexcitability (5) that affects neuronal network circuitry. However, as glial cells are integral players in synaptic function and neuronal circuits, their malfunction can also promote neuronal hyperexcitability and hypersynchrony (6–9). Surprisingly, despite the known bi-directional communication between glia and neurons, our understanding of the involvement of glial cells in AD pathology, specifically their role in mediating neuronal signalling, remains limited.

Astrocytes, a subtype of glial cells, are actively involved in the modulation of neuronal activity, in addition to their more established supportive functions in transporting metabolites and maintaining the homeostasis of the extracellular matrix (10). One of the prominent features of AD is the transition of astrocytes into a reactive state (“*reactive astrocytes*”) throughout disease progression, affecting both their function and morphology (11, 12). Indeed, mounting evidence from human patients and mouse models suggests neuroinflammation, in which astrocytes and microglia become more reactive, is an active player contributing to the pathogenesis of AD, reviewed by (13).

The maintenance of ion concentrations in the extracellular milieu is essential to preserve transmembrane ion gradients and thus vital for many physiological processes, including the initiation of action potentials, enhancement of synaptic plasticity and rhythmogenesis (14–17). Astrocytes play a key role in maintaining ion homeostasis in the extracellular space, and it has been suggested that dysregulation of ion homeostasis can affect neuronal function (7, 18, 19) and serve as a trigger to neurodegeneration, reviewed by (20).

While the majority of AD cases are sporadic (sAD), about 5% of all AD patients carry a familial mutation (fAD) either in the amyloid precursor protein (APP), presenilin-1 or presenilin-2 (21). These hereditary mutations led to the development of animal models, such as the 5xFAD transgenic mouse that expresses five familial AD mutations (APP KM670/671NL (Swedish), APP I716V (Florida), APP V717I (London), PSEN1 M146L, PSEN1 L286V) under the control of the neuronal promotor Thy1, that additively increase Aβ production (22). The 5xFAD mice develop robust amyloid pathologies, with plaques accumulating in the brain from 2 months of age, triggering microgliosis and inflammatory processes, as well as synaptic and neuronal loss (23). Similar to AD patients, the 5xFAD mice developed age-related impairments in many forms of learning and memory, including cued and/or contextual associative fear, object and spatial recognition, conditioned taste aversion and visuo-spatial navigation and reference memory (24).

We previously showed that the excitability and oscillatory profile of individual neurons as well as neuronal networks changes following modulation of the astrocytic [K^+^]_o_ clearance (7). Hence, this study aims to investigate the ability of the panglial syncytium to clear [K^+^]_o_ from the extracellular matrix in the brain of 5xFAD mice brains during disease progression. To this end, we compared the functional and anatomical features of astrocytes during disease progression at two brain regions associated with AD, the somatosensory cortex and hippocampus. We further assessed the activity of nearby neurons and compared them to age-matched wild-type littermates to better understand the impact of astrocytic dysfunction on neuronal signalling.

## Materials and Methods

### Animals

Transgenic 5xFAD mice (#034840, Jackson Laboratories) were housed under standard conditions of temperature, humidity, 12h light/dark cycle, free access to food and water, and without any intended stress stimuli. All experiments were approved and performed in accordance with Western Sydney University committee for animal use and care guidelines (Animal Research Authority #A14455).

### Slice preparation

Mice were deeply anesthetized by inhalation of isoflurane (5%) and then injected with a ketamine/xylazine cocktail (100mg/kg and 200mg/kg, respectively) as previously described (25, 26). Following anaesthesia, mice were transcardially perfused with ice-cold protective recovery artificial CSF (aCSF) containing (in mM): 92 mM NMDG, 2.5 KCl, 1 MgSO_4_, 1.25 NaH_2_PO_4_, 2 CaCl_2_, 30 NaHCO_3_, 20 HEPES, 25 glucose, 2 Thiourea, 5 Na-ascorbate, 3 Na-pyruvate and saturated with carbogen (95% O_2_−5% CO_2_ mixture; pH 7.4), until the outflow solution was clear. Following perfusion, the mice were decapitated, the brains were quickly removed and placed into ice-cold protective recovery aCSF (as above). Coronal brain slices (300μm thick) were cut with a vibrating microtome (Leica VT1200S) and transferred to the Braincubator^TM^ (PaYo Scientific, Sydney; http://braincubator.com.au) with incubation aCSF containing (in mM): 125 NaCl, 2.5 KCl, 1 MgCl_2_, 1.25 NaH_2_PO_4_, 2 CaCl_2_, 25 NaHCO_3_, 25 glucose and saturated with carbogen (95% O_2_−5% CO_2_ mixture; pH 7.4), as reported previously (27, 28). The Braincubator^TM^ is an incubation system that closely monitors and controls pH, carbogen flow, and temperature, as well as irradiating bacteria through a separate UV chamber (29, 30). Slices were initially incubated for 12 min at 30^◦^C, after which they were incubated with sulforhodamine 101 (SR101, 1μM at 37^°^C for 20 minutes), a water-soluble red fluorescent dye used for labelling astrocytes and oligodendrocytes (31), and then cooled to 15–16^◦^C and kept in the Braincubator^TM^ for at least 30 min before measurements were taken (32).

### Electrophysiological recordings and stimulation

The recording chamber was mounted on an Olympus BX-51 microscope equipped with infrared/differential interface contrast (IR/DIC) optics. Following incubation in the Braincubator^TM^, slices were transferred to the recording chamber and left for a minimum of 5 minutes to allow them to be warmed to 36^°^C. Slices were constantly perfused with carbogenated aCSF at a rate of 2 ml/min.

Whole-cell recordings from astrocytes and neurons in L2/3 of the somatosensory cortex (SSC) and stratum lacunosum moleculare (SLM) of CA1 region of the hippocampus were obtained with patch pipettes with a typical resistance of 5–7 MΩ containing (in mM): 126 K^+^-gluconate, 4 KCl, 10 HEPES, 0.3 EGTA, 0.3 Na_2_GTP, 2 Na_2_ATP, 4 MgATP, 10 phosphocreatine, pH adjusted to 7.3 (285-290 mOsm). Voltages were recorded in current clamp mode using a Multiclamp 700B dual patch-clamp amplifier (Axon Instruments, Foster city, CA), digitally sampled at 30 – 50kHz, filtered at 10kHz, and analysed off-line using pCLAMP10. The series resistance and capacitance (up to 80%) were compensated automatically to minimize voltage errors.

For measurement of Ba^2+^ sensitive currents, astrocytes were voltage clamped at -70 mV and then stepped to command voltages (1 sec) from -200 mV to +70 mV with 20 mV increments. The Kir component was isolated by subtracting the currents following Ba^2+^ application from the baseline currents measured before the application of Ba^2+^ (100μM). Cells were considered stable and suitable for analysis if the access resistance, input resistance and resting membrane potential did not change by more than 20% from their initial value during recording.

### Measuring K^+^ clearance rate

Preparation and calibration of the K^+^-selective microelectrodes were performed as per (33). In short, the voltage response of the salinized K^+^-selective microelectrodes was calibrated before and after experiments within the experimental chamber by placing the electrode in aCSF containing different KCl concentrations. Once the electrode potential reached a steady state, a dose-response curve was calculated using a half-logarithmic (Log_10_) plot. Only electrodes showing slopes of 53-60 mV for a 10-fold change in [K^+^]_o_ were used. K^+^-selective microelectrodes were considered good if the recorded voltage baseline was stable and the voltage response was similar before and after its experimental usage (∼20% deviation). Various KCl concentrations were applied to brain slices at a constant distance of ∼10 μm from the K^+^-selective microelectrode through a puffing pipette (tip diameter of 1μm) for 0.1 seconds at 10 psi, as previously described (34, 35). The rate constant (λ = 1/τ) and decay time (defined as the time it took the amplitude to decay from 90-10%) were used for statistical analysis. K^+^ “clearance rate” and “rate constant” are used interchangeably.

### Assessment of the astrocytic syncytium

To assess astrocytic coupling, individual astrocytes were loaded with an intracellular solution containing biocytin (0.3%; Sigma) for 12 min in a whole-cell mode, as previously described (7, 34, 35). Hyperpolarizing step currents (50 pA for 500 ms at 0.5 Hz) were applied via the recording electrode to facilitate the spreading of the biocytin to neighbouring astrocytes, as previously described.(36, 37) The slice was then fixed in paraformaldehyde solution (PFA, 4%) for < 24 hours. Following washing in PBS to clear the PFA, slices were incubated with Alexa Fluo-488-conjugated streptavidin (1:200, Thermo Fischer, #S11223) dissolved in PBS and 0.1% Triton-X for 72 hrs at 4°C. Slices were washed with PBS, mounted onto glass slides with Vectashield fluorescent mounting medium (Vector Laboratories #H-1200), and dried overnight. Images of astrocytic morphology were taken with a confocal microscope (Zeiss, LSM 5 Pascal).

Stained astrocytes were counted using the region of interest function in FIJI software. In short, the average intensity of biocytin signals was used to distinguish recorded cells from other cells that were coupled to it. If the mean fluorescence intensity of a connected astrocyte was above 75% of the recorded astrocyte, it was considered to be a primary connection. For each slice, z-stacks were acquired to include the entire biocytin-stained syncytia.

### Histology and Immunohistochemistry

Mice brains were quickly removed and placed in a 4% PFA solution for at least 24 hrs followed by cryoprotection in increasing concentrations of sucrose solution (10%, 20% and 30% in PBS) for at least 48 hrs per concentration. 50μm coronal sections were cut with a Leica cryostat (Leica CM1950). The slices were washed in PBS and then frozen in cryoprotectant solution (30% ethylene glycol, 30% glycerol in PBS) until staining.

To determine the expression of astrocytic Kir4.1 channels, slices were incubated with GFAP and Kir4.1 antibodies. Prior to incubation with the antibodies, slices were washed in PBS (x3) to remove any remaining cryoprotectant solution and subjected to heat-induced antigen retrieval in 10 mM sodium citrate (pH=6) for 30 mins at 80°C. Next, slices were washed in PBS and incubated in a blocking solution (normal donkey serum 5% with Triton-X 0.1% in PBS) for 2 hrs at room temperature. Following washing in PBS, slices were incubated with Kir4.1-antibody (rabbit-anti-Kir4.1; 1:200; Alomone Labs, APC-035) and GFAP antibodies (GFAP clone antibodies #MAB360; 1:200, Millipore) for 24 hrs at 4°C. Next, slices were washed with PBS and incubated with fluorescent secondary antibodies (Alexa Fluor 594-AffiniPure Donkey Anti-Rabbit; Alexa Fluor 488-AffiniPure Donkey Anti-mouse; Jackson Labs; 1:200) with 0.1% Triton-X in PBS for 2 hrs, followed by washing in PBS. The slices were transferred onto glass slides and mounted with DAPI-containing Vectashield fluorescent mounting medium until analysis with a confocal microscope (Zeiss, LSM 800) at 488nm and 594nm. Images of astrocytes were acquired using high (x63) magnification objective lenses with 0.4 µm step for 3D reconstruction.

### Astrocytic 3D reconstruction

Reactive astrocytes are characterised by enhanced expression of GFAP as well as structural changes that affect their morphology, including cellular hypertrophy, elongation of astrocytic processes, and overall increase in the cell volume (38). To assess astrocytic reactivity, GFAP stained astrocytes were reconstructed in 3D with Neurolucida 360 software (MBF Bioscience) and analysed with a built-in analysis software (Neurolucida Explorer) to determine morphological features such as the soma size and volume, as well as the overall length and thickness of the astrocytic processes.

### Western Blot analysis

To investigate protein expression, hippocampus and somatosensory cortex samples were micro dissected from whole brains, snap-frozen in liquid nitrogen and stored at –80 °C until ready for use. Then, tissue samples were homogenized in homogenisation buffer (8 M urea, 2 M thiourea, 4% CHAPS at 4 °C) containing protease phosphatase inhibitors and 1 mM sodium orthovanadate and sonicated with a probe sonicator (VTSYIQI Integrated Ultrasonic Homogenizer Sonicator, 50W for 2 s with an interval of 10 s between the sonication cycles, 3-5 cycles). Following homogenisation, the sample was transferred to ultracentrifuge tubes (Ultra Clear Centrifuge Tubes, Beckman Cat # 344057) and centrifuged at 50000 g (SW 55 Ti rotor, Beckman Coulter, Indianapolis, IN, USA) at 4°C for 1 h. The supernatants were collected into individual tubes for either immediate analysis or storage at −80 ^0^C. The protein concentration was estimated using the EZQ™ protein quantitation kit (Life Technologies, Eugene, OR, USA) according to the manufacturer’s instructions. Equal amounts of protein were boiled for 3 min with Laemmli-SDS sample buffer (63 mM Tris HCl, 20% Glycerol, 2% SDS, 0.0025% Bromophenol Blue, pH 6.8) containing 2.5% b-mercaptoethanol. After the cooling, 20 µg of total proteins were run on a 10% SDS –polyacrylamide gel and separated for 3 hours at 4 °C (10 min at 120V followed by 170 minutes at 90V). Separated proteins were transferred onto a polyvinyl difluoride (PVDF) membrane (0.45μm; Thermofisher) for 22 hours at 4°C (30 mA per gel). Then, the membrane was blocked overnight at 4 °C in 5% non-fat dry milk dissolved in Tris-buffered saline containing 0.1% Tween (TBST) and 3% polyvinylpyrrolidone (PVP). Following blocking, the membrane was washed for 50 minutes with TBST buffer (5x10 minute washes). To probe with the membrane, primary anti-rabbit Kir4.1 antibodies (Alomone; 1:500) were diluted in TBST buffer with 3% PVP and incubated with the membrane for 24 hours at 4°C. Following incubation, the membrane was washed with TBST buffer and the membrane was set for 2 hours at room temperature with the secondary goat anti-rabbit IgG HRP antibodies (Sigma Aldrich; 1:10000) diluted in a TBST buffer with 3% PVP. Blots were visualised using Luminata™ Forte western HRP substrate (Millipore, WBLUR0100, 4 min exposure). To normalise the signal, the expression of ß-actin was analysed in the same blot after stripping (200mM NaOH, 20 min) using anti-ß-actin antibodies (Sigma-Aldrich; 1:500). Blots were visualised using Luminata™ Crescendo western HRP substrate (Millipore, WBLUR0100, 1 min exposure). All Western blot experiments were repeated five times and analysed using Image Lab Software 6.1 (Bio-Rad).

### Experimental design and statistical analysis

Detailed experimental designs of K^+^ clearance measurements are described in the results sections. Data were assessed for normality and variance homogeneity using the D’Agostino & Pearson test and Bartlett’s test, respectively. Statistical comparisons were done using two-way or three-way ANOVA followed by Sidak’s post hoc test, according to the experimental design. Statistical comparisons were performed with Prism 10 (GraphPad Software; San Diego, CA, USA), and unless stated otherwise, data is reported as mean ± S.E.M. Probability values <0.05 were considered statistically significant.

## Results

K^+^ homeostasis is highly dependent on the ability of astrocytes to clear K^+^ from the extracellular milieu. To measure the clearance rate during the progression of AD, we have recorded the [K^+^]_o_ in the hippocampus and somatosensory cortex of transgenic mice expressing 5 different familial mutations (5xFAD mice) at different disease stages. 5xFAD mice express several symptomatic hallmarks that are associated with AD, including a significant build-up of amyloid plaques(22) (see also Fig. 1), neuronal loss, enhanced neuroinflammation and synaptic deficits (39). Based on the expression of these symptoms, we have measured the [K^+^]_o_ clearance rate in 5xFAD mice during their “presymptomatic” phase (6-8 weeks, termed ‘Young 5xFAD’, *N*=49), while no *β*-Amyloid aggregates were noticeable, and at their “symptomatic” stage (26-30 weeks, termed ‘Aged 5xFAD’, *N*=75), while both the distribution of *β*-Amyloid plaques and neuroinflammation levels were very high (Fig. 1A-F). As controls, we used age-matched WT littermates; hence, we named them “Young WT” (*N*=48) and “Aged WT” (*N*=72), respectively.

**Figure 1.**
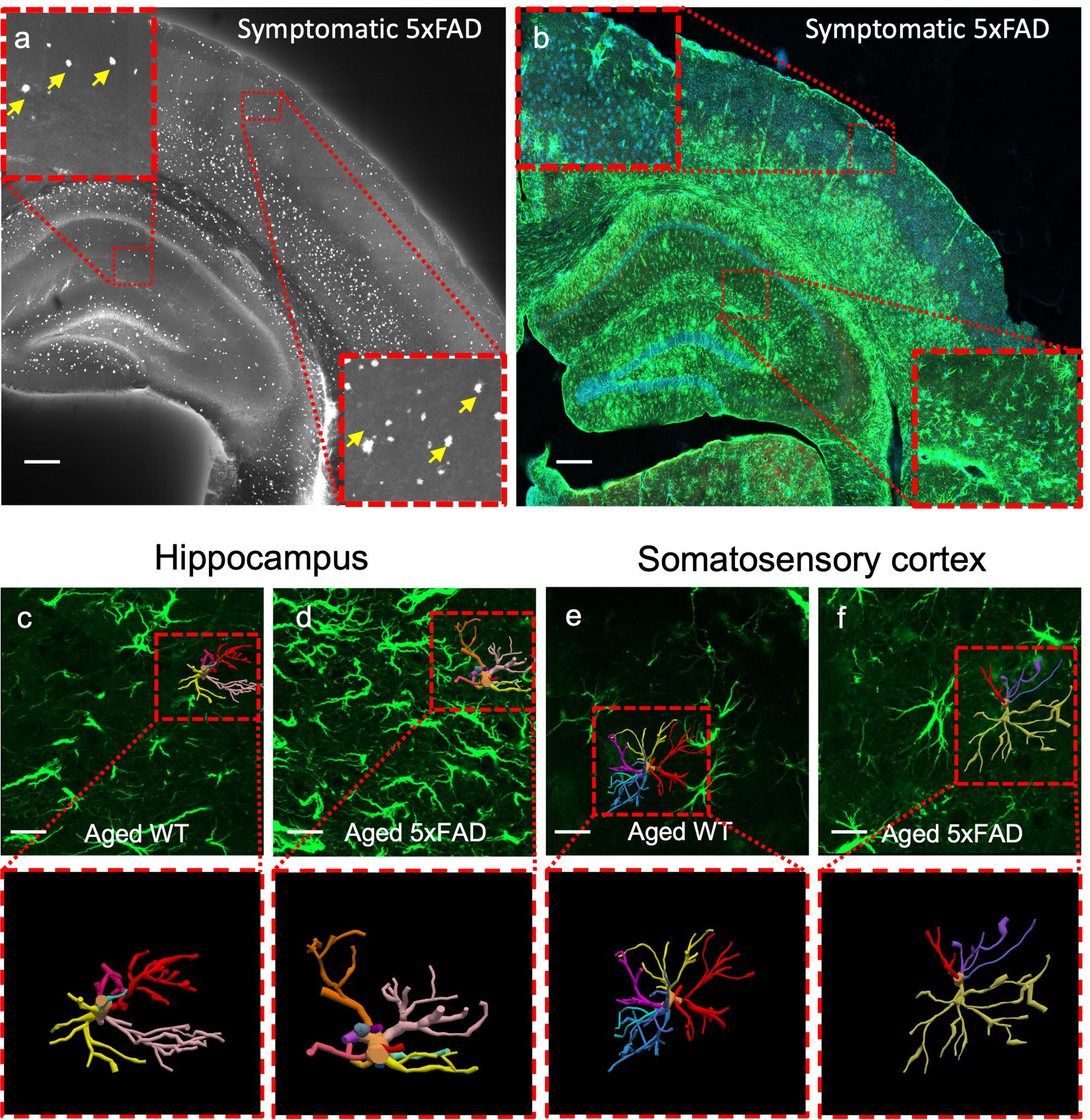
Amyloid pathology in aged 5xFAD mice. A-B) Confocal images depicting A) B-amyloid plaques (yellow arrows) labelled with Thioflavin S in the hippocampus and somatosensory cortex of aged (6 months) 5xFAD transgenic mice. Insets – high magnification image depicting the ß-amyloid plaques in the somatosensory and SLM of CA1. B) The amyloid accumulation was associated with strong astrogliosis depicted by GFAP immunostaining in aged 5xFAD mice. GFAP staining in green, DAPI in blue. C-F) Confocal images depicting astrocytic morphology in the hippocampus (C-D) and somatosensory cortex (E-F) of aged WT (C, E) and aged 5xFAD (D, F) mice. Inset – 3D reconstruction of astrocytes.

### Increased astrocytic reactivity in the hippocampus of symptomatic 5xFAD mice

Previous reports indicated that AD is associated with astrocytic reactivity, characterised by morphological changes and increased expression of the glial fibrillary acidic protein (GFAP) (38, 40). To assess the morphological features of astrocytes in 5xFAD mice, we analysed their 3D morphology in the hippocampus and somatosensory cortex of 5xFAD and WT littermates (Fig. 1-2). Reconstructed astrocytes from the hippocampus of aged 5xFAD mice indicated significant hypertrophy compared to age-matched control (Fig. 1C, D), while astrocytes that reside in the somatosensory cortex were comparable between groups (Fig. 1E, F). Moreover, the average soma volume, soma diameter, processes length and processes thickness of **hippocampal astrocytes** were significantly higher in aged 5xFAD mice than in their age-matched controls, and these changes were due to the familial mutations and not due to ageing (two-way ANOVA, Fig. 2). Indeed, while all parameters of astrocytic morphology were comparable between young 5xFAD mice and their age-matched littermate controls (Fig. 2), the average soma volume in the hippocampus of aged 5xFAD mice was 40.25 ± 8.71 µm^3^ (*n* = 10) compared to 10.40 ± 1.52 µm^3^ (*n* = 9) in age-matched WT (*p* <0.001, two-way ANOVA with Sidak’s post hoc test, Fig. 2A). The soma diameter of astrocytes in aged 5xFAD mice was higher by 44% then in aged WT mice (3.92 ± 0.34 µm, *n* = 10 vs. 2.73 ± 0.18 µm, *n* = 9, respectively, *p* =0.007, two-way ANOVA with Sidak’s post hoc test, Fig. 2B). Also, the length and the thickness of astrocytic processes from aged 5xFAD mice were higher by 72% and 18% respectively (processes length: 381.02 ± 57.18 µm, *n* = 9 vs. 220.21 ± 32.1 µm, *n* = 9, *p* <0.01; processes thickness: 4.54 ± 0.16 µm, *n* = 9 vs. 3.82 ± 0.14 µm, *n* = 9, *p* <0.01, two-way ANOVA with Sidak’s post hoc test, Fig. 2C, D). In contrast, astrocytic morphology in the **somatosensory cortex** was largely comparable between all groups (Fig. 2E-F), indicating that astrocytic reactivity was region-specific and not uniform across all brain regions. We next assessed astrocytic connectivity via intracellular injections of 0.3% biocytin and evaluation of the number of primary coupled astrocytes to the patched astrocytes as previously described (7, 34, 35). Our data indicate a significant decrease in the number of primary coupled astrocytes in the hippocampus of 5xFAD mice, which was due to the 5xFAD mutations (*F*(_1, 15_)=7.33, *p*<0.01 for the factor ‘*5xFAD mutations’*; *F*(_1, 15_) = 0.14, p=0.7 for the factor ‘*aging*’ and *F*(_1, 15_) = 1.57, *p* = 0.22 for the factor ‘*Interaction*’, two-way ANOVA). On average, the number of primary connected astrocytes in the hippocampus of aged 5xFAD mice was significantly lower than in their age-matched controls (2.75±1.03, *n* = 5 vs. 8.2±2.02, *n* = 5, respectively, *p* = 0.03, two-way ANOVA with Sidak’s post hoc test, Fig. 2K). In contrast, astrocytic connectivity at the somatosensory cortex was comparable between groups (two-way ANOVA, Fig. 2I-J), further indicating on region specific changes in astrocytic reactivity.

**Figure 2.**
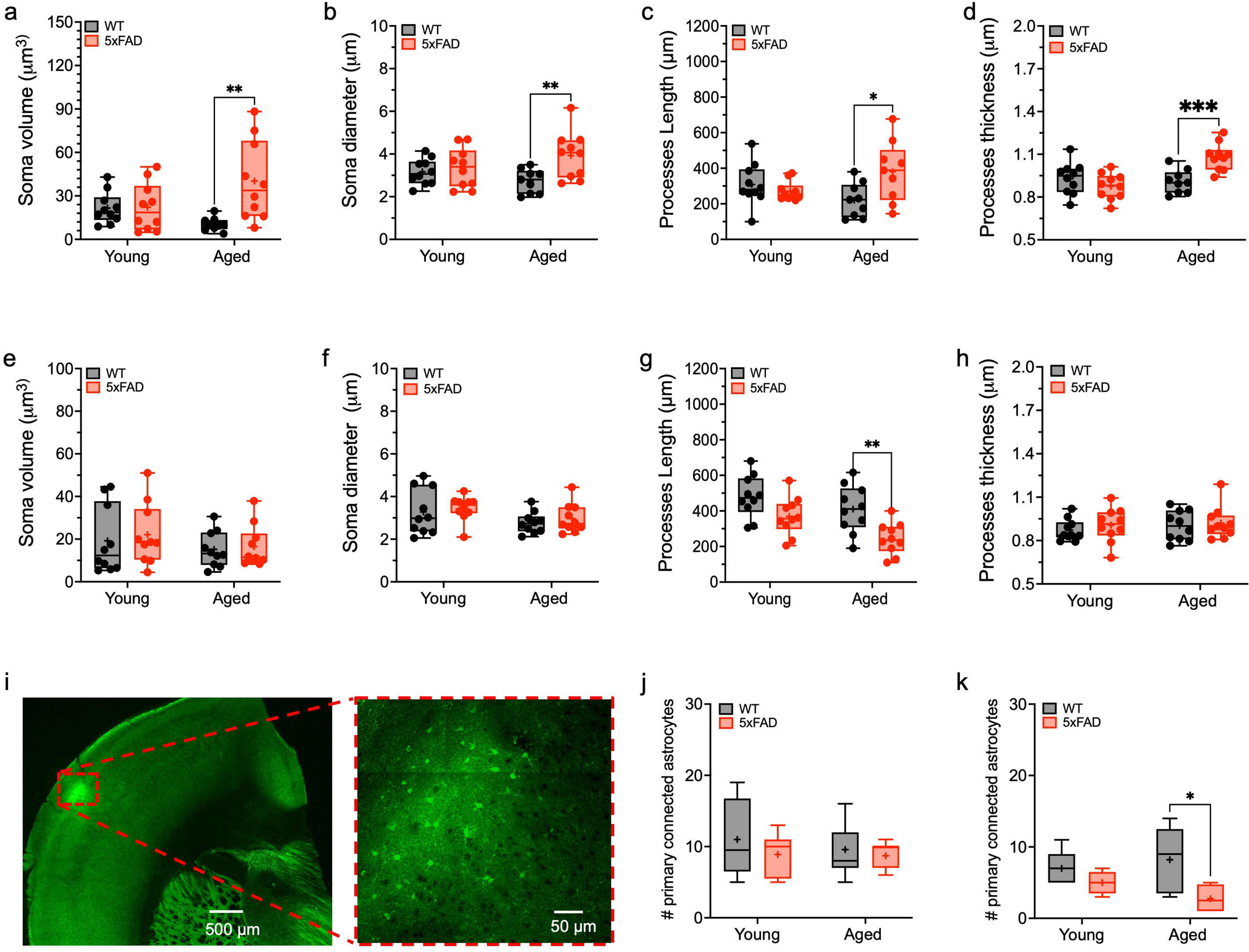
Reactive astrocytes in the hippocampus of 5xFAD mice. Box plots depicting astrocytic morphology in the hippocampus (A-D) and somatosensory cortex (E-H) of all mice, including the average soma volume (A, E), diameter (B, F), processes length (C, G) and processes diameter (D, H). I-L) Confocal images of biocytin stained astrocytes in the hippocampus (I) and somatosensory cortex (K). Insets – high magnification (x63) of the delimited areas. J, L) Box plots depicting the number of primary connected astrocytes in the hippocampus (J) and the somatosensory cortex (L). The box upper and lower limits are the 25th and 75th quartiles, respectively. The whiskers depict the lowest and highest data points, the horizontal line through the box is the median, and the “+” sign represents the mean. *p < .05, **p < .01, ***p < .001; ****p < .0001; two-way ANOVA with Sidek’s post hoc test.

### Region-specific impairment of [K^+^]_o_ clearance rate

K^+^ clearance rate was assessed via local measurements of the extracellular K^+^ concentration ([K^+^]_o_) prior to and following transient local application of KCl at 3 different concentrations corresponding to *low*, *high* and *excessive* [K^+^]_o_ (5 mM, 15 mM and 30 mM respectively; Fig. 3A, B), as previously described (34). These concentrations were elected as they are correlated to different levels of network activity and involves different phases within the astrocytic clearance mechanisms (7, 8). The [K^+^]_o_ clearance rate constant (λ = 1/τ) was calculated from the fitted exponential decline of the voltage signal (90%–10%) using a least-squares fitting routine (Chebyshev algorithm Clampfit 10, Molecular devices), as previously described (35).

**Figure 3.**
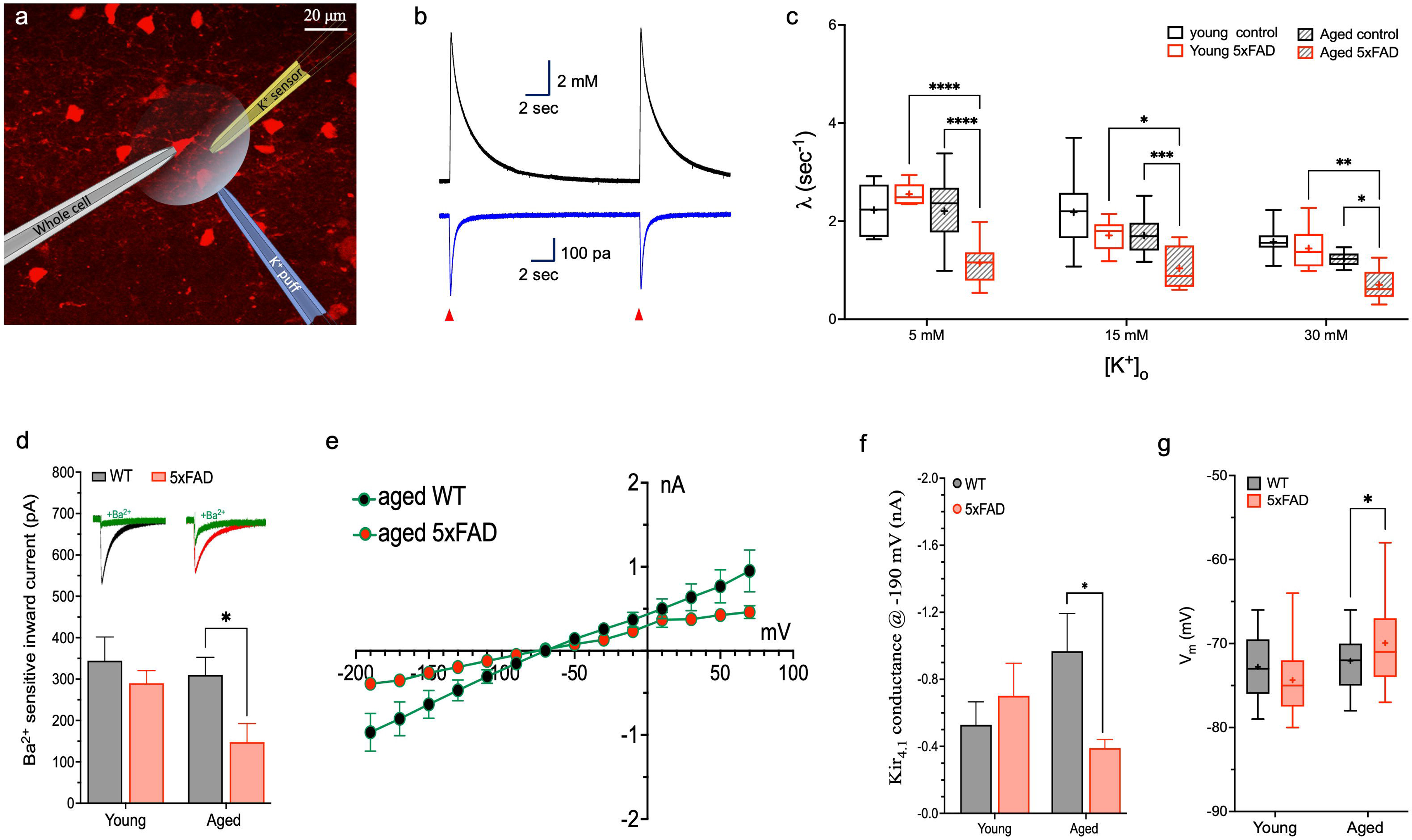
Astrocytic K^+^ clearance in the hippocampus of 5xFAD mice. A) Fluorescence image of a brain slice depicting astrocytes labelled with SR101 (red) along with a schematic diagram illustrating the experimental design, including the K^+^ selective recording electrodes (K^+^ sensor), K^+^ puffing pipette (K^+^ puff) and the whole-cell recording electrode (whole-cell). (B) Sample traces of extracellular [K^+^]_O_ recorded with the K^+^ selective electrode (top) and intracellular influx current into nearby astrocytes depicting the K^+^ clearance time course following local application of 30 mM. C) Box plot depicting the clearance rate in the hippocampus of young and aged mice following application of KCl at different concentrations. The box upper and lower limits are the 25th and 75th quartiles, respectively. The whiskers depict the lowest and highest data points, the horizontal line through the box is the median and the “+” sign represents the mean. \**p* < .05, \*\**p* < .01, \*\*\**p* < .001; \*\*\*\**p* < .0001; three-way ANOVA with Tukey’s post hoc test. D) Bar graph depicting the average Ba^2+^ sensitive current following application of 30 mM KCl Inset – sample traces taken from aged WT (black) and aged 5xFAD (red) mice before and following Ba^2+^ application (green). E) I–V plot and bar graph (F) showing the average Ba2+ sensitive currents in the hippocampus of aged 5xFAD mice and their age-matched littermate controls. \**p* < 0.05, student t-test.

To assess the impact of ‘*ageing*’, ‘*5xFAD mutations*’ and ‘*K^+^ concentration*’ on the clearance rate, we performed a three-way ANOVA analysis on these variables. Our results indicate that the clearance rate recorded in the hippocampus was significantly affected by all three factors (*F*(_1, 149_) = 70.89, *p* < 0.0001 for the factor ‘*aging*’; *F*(_1, 149_) = 34.17, *p* < 0.0001 for the factor ‘*5xFAD mutations’* and *F*(_2, 149_) = 39.25, *p* < 0.001 for the factor ‘*K^+^ concentration’*), with significant two-way interaction between ‘*ageing x 5xFAD mutations’* (*F*(_1, 149_) = 20.62, *p* < 0.001), and three-way interaction between all factors (*F*(_2, 149_) = 6.266, *p* < 0.005, three-way ANOVA, Figure 3C). Post hoc analysis suggested that following local transient application of low concentration of KCl (5 mM) the average clearance rate constant (λ) was comparable between young WT and young 5xFAD mice, but was significantly lower in the hippocampus of aged 5xFAD mice compared to their aged-match WT (1.14±0.1 s^-1^, *n* =16 vs 2.20±0.16 s^-1^, *n* = 17, *P* < 0.0001, three-way ANOVA with Tukey’s post hoc test), or young 5xFAD mice (*P* < 0.0001, three-way ANOVA with Tukey’s post hoc test, Fig. 3C). Similarly, the K^+^ clearance rate following transient local application of high (15 mM) or excessive (30 mM) concentrations of KCl was significantly lower in the aged 5xFAD mice compared to their age-match WT (1.03±0.1 s^-1^, *n* = 13 vs. 1.7±0.07 s^-1^, *n* = 24, P<0.0001 at 15 mM and 0.7±0.09 s^-1^, *n* = 12 vs. 1.2±0.04 s^-1^, *n* = 8, *p* < 0.05 at 30 mM, Tukey’s post hoc test) or young 5xFAD mice (1.7±0.08 s^-1^, *n* = 11, P<0.01 at 15 mM and 1.4±0.1 s^-1^, *n* = 13, *p* < 0.001 at 30 mM, Tukey’s post hoc test, Fig. 3C). These results suggest that the astrocytic K^+^ clearance rate in the hippocampus of 5xFAD mice is altered during disease progression.

To assess the mechanisms underlying the altered clearance rate, we next investigated specific cellular and intracellular processes that impact the clearance rate, including K^+^ uptake mediated via inward rectifying K^+^ (Kir) channels and K^+^ spatial buffering via the astrocytic syncytium (8, 41, 42). Within the K^+^ inward rectifying channel family, Kir_4.1_ channels are specifically expressed in astrocytes and oligodendrocytes and are responsible for the majority of K^+^ uptake following neuronal activity, reviewed by (35, 43, 44). Currently, there is no specific blocker for Kir4.1 channels, however previous studies have shown that a low concentration of BaCl_2_ (100 μM) can selectively block Kir_4.1_ channels and thus impair the K^+^ uptake mechanism in astrocytes (7, 34, 37, 45–47). Indeed, the clearance rate recorded in the hippocampus following thevapplication of Ba^2+^ (100 μM for 5 min) indicated a significant decrease of the clearance rate constant in all groups, except for the aged 5xFAD mice group, in which λ was comparable across all recorded [K^+^]_o_ concentrations (Table 1).

**Table 1.** The impact of Kir4.1 channels blockade on the clearance rate constant (λ). The impact of Kir4.1 channels blockade on the K^+^ clearance rate constant at different [K^+^]_O_ concentrations. The % of change was determined by subtracting the clearance rate following application of Ba^+2^ [K_Ba_^2+^] from the baseline clearance rate (K_baseline_ measured before Ba^2+^ application) and dividing it by the baseline clearance rate (K_baseline_). Paired t-test was used to tease out the impact Kir4.1 blockade in each concentration. Significance levels presented as *p < 0.05, **p < 0.005.

| Brain region |  | Hippocampus |  |  |  | Somatosensory cortex |  |  |  |
| --- | --- | --- | --- | --- | --- | --- | --- | --- | --- |
| Experimental group |  | *Young control*<br>*n=30* | *Young 5xFAD*<br>*n=26* | *Aged control*<br>*n=30* | *Aged 5xFAD*<br>*n=38* | *Young control*<br>*n=31* | *Young 5xFAD*<br>*n=29* | *Aged control*<br>*n=29* | *Aged 5xFAD*<br>*n=36* |
| Clearance rate with Ba<sup>2+</sup> (% from baseline) | **5 mM** | -30±7* | -20±8* | -22±5** | -1±5 | -18±7* | -22±9* | -21±5* | -14±3** |
|  | **15 mM** | -40±6** | -8±2* | -23±7* | -1±5 | -12±3* | -24±8* | -24±9* | -16±4** |
|  | **30 mM** | -32±8** | -11±4* | -36±19* | -2±6 | -16±6* | -20±7* | -15±4* | -23±6** |

We next used whole-cell voltage clamp recording from astrocytes prior to and following bath application of Ba^2+^ to assess the overall conductance via Kir_4.1_ channels (48). Ba^2+^ sensitive inward currents were determined by subtracting the inward current recorded following the application of Ba^2+^ from the baseline current (Fig. 3D). Comparison of Ba^2+^ sensitive inward currents in hippocampal astrocytes which were recorded during transient application of 30 mM KCl indicated a significant decrease in aged 5xFAD mice compared to their age-matched controls (*F*(_1, 19_) = 3.83, *p* < 0.065 for the factor ‘*aging*’; *F*(_1, 19_) = 5.8, *p* < 0.02 for the factor ‘*5xFAD mutations’* and *F*(_1, 19_) = 1.43, *p* < 0.24 for the factor ‘Interaction’, Two-way ANOVA). While the average inward current was comparable between young control, young 5xFAD and aged control (344 pA (*n* = 6), 310±57 pA (*n* = 6) and 289±30 pA (*n* = 6), respectively), it significantly decreased in aged 5xFAD mice (147±44 pA, *n* = 6, *p* < 0.05, Tukey’s post hoc test Fig.3D). Moreover, the overall Kir_4.1_ conductance in hippocampal astrocytes of aged 5xFAD mice was significantly lower than in their age-matched control (Fig. 3E-F), indicating altered functionality that occurs during disease progression and is independent of the extracellular K^+^ concentration. In line with these results, astrocytic resting membrane potential (V_m_) was found to be more depolarized in the hippocampus of aged 5xFAD mice (two-way ANOVA, Fig. 3G), consistent with previous studies (49, 50). These results were specific to the hippocampus, as measurement of the clearance rate, as well as Ba^2+^ sensitive currents at the somatosensory cortex indicate comparable resultes between all groups and across all concentrations (three-way ANOVA, Fig. 4).

**Figure 4.**
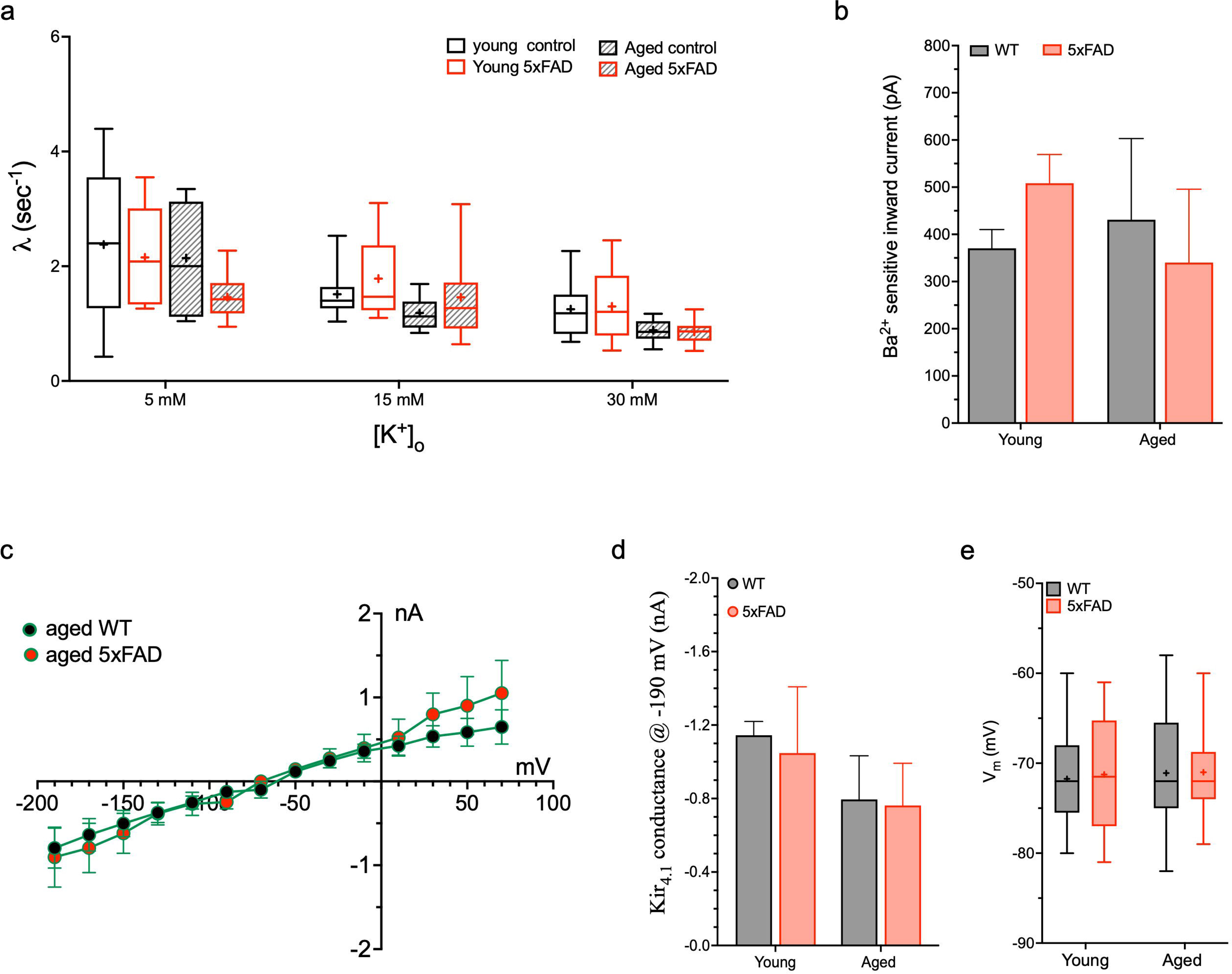
Astrocytic K+ clearance in the somatosensory cortex of 5xFAD mice. A) Box plot depicting the clearance rate in the somatosensory cortex of young and aged mice following application of KCl at different concentrations. The box upper and lower limits are the 25th and 75th quartiles, respectively. The whiskers depict the lowest and highest data points, the horizontal line through the box is the median, and the “+” sign represents the mean (three-way ANOVA). B) Bar graph depicting the average Ba^2+^ sensitive current following application of 30 mM KCl. C) I–V plot and bar graph (D) showing the average Ba2+ sensitive currents in the somatosensory cortex of aged 5xFAD mice and their age-matched littermate controls.

To further investigate the causes underlying the decrease in Kir4.1 conductance in aged 5xFAD mice, we have measured the overall expression of Kir4.1 channels in both hippocampus and somatosensory cortex (Fig. 5). Kir4.1 protein can be expressed as a glycosylated (50 kDa) or aglycosylated protein (37 kDa) (51). However, while in astrocytes most Kir4.1 channels are highly glycosylated, oligodendrocytes predominantly express the aglycosylated form of Kir4.1 channels (52). Analysis of the expression level of glycosylated (50 kDa) kir4.1 proteins indicated a significant decrease in the hippocampus of aged 5xFAD mice which was affected by both 5xFAD mutations and ageing (*F*(*_1, 8_*) = 5.39, p=0.04 for the factor ‘aging’; *F*(*_1, 8_*) = 8.89, p=0.01 for the factor ‘5xFAD mutations’ and *F*(*_1, 8_*) = 3.36, *p* = 0.1 for the factor ‘Interaction’, Two-way ANOVA). Indeed, the average expression level of 50 kDa Kir4.1 was significantly lower in the hippocampus of aged 5xFAD mice vs age-matched controls (1.74 ± 0.3, *n* = 5, vs 3.78 ± 0.5, *n* = 5, respectively, *p* <0.01, two-way ANOVA with Sidak’s post hoc test, Fig. 5C). In contrast, the expression level of (50 kDa) kir4.1 proteins in the somatosensory cortex was comparable between groups (Fig. 5D). Further analysis of the aglycosylated form of Kir4.1 channels (37 kDa) indicated comparable results between all groups tested, at both hippocampus and somatosensory cortex (Fig. 5E-F). These results suggest that the expression level of Kir4.1 channels is reduced specifically in astrocytes residing in the hippocampus.

**Figure 5.**
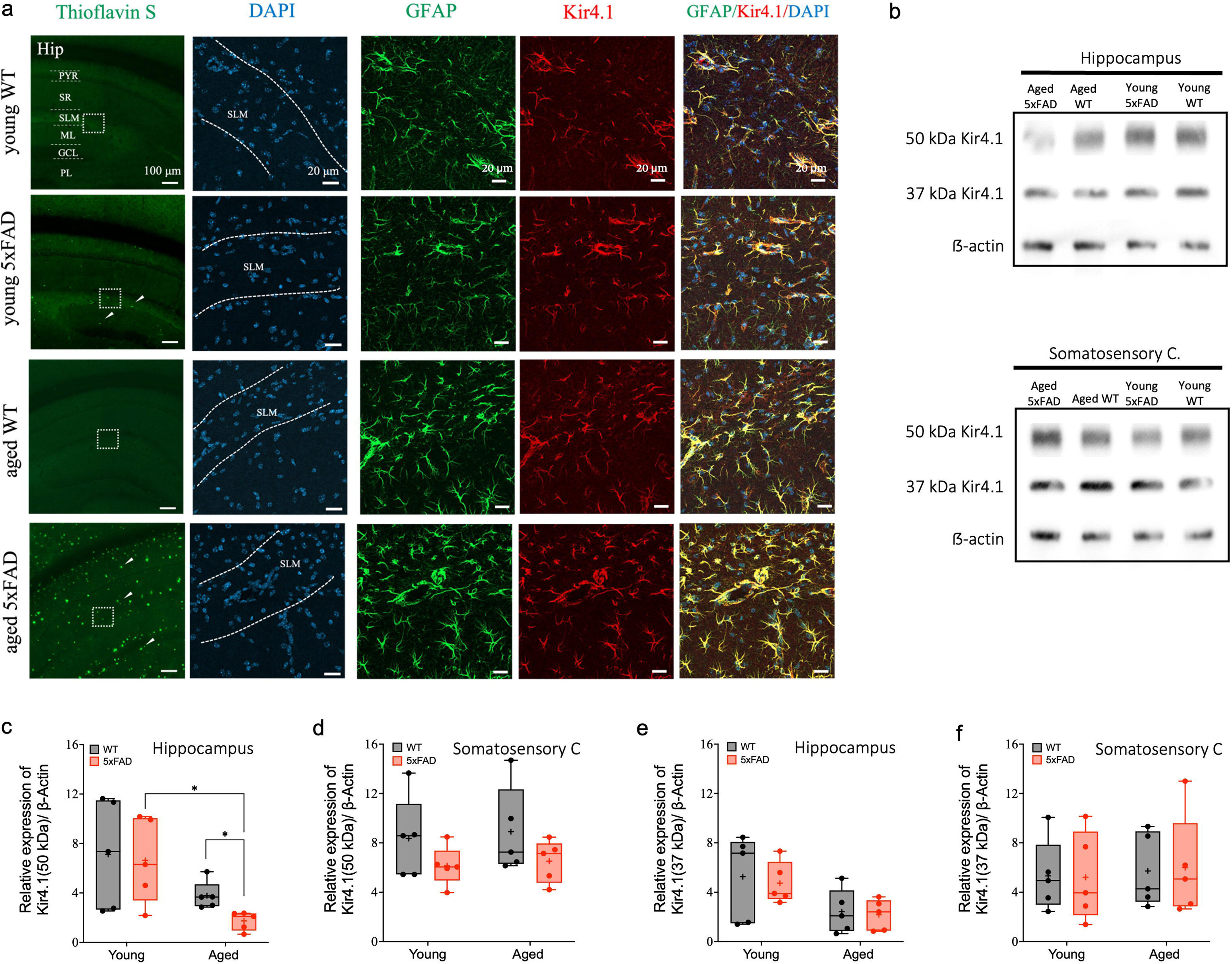
Expression of Kir4.1 channels in the hippocampus and somatosensory cortex of 5xFAD mice. A) Left – Low magnification (x10) confocal images depicting *Aβ plaques (arrows) labelled* with Thioflavin-S staining (green). High magnification (x40) staining of the SLM region (marked with dashed lines) depicting cells nuclei (DAPI), astrocytes (GFAP) and Kir4.1 (red) staining in the hippocampus of 5xFAD mice and their age-matched littermate controls. *PYR - pyramidal cell layer, SR - stratum radiatum, SLM - stratum lacunosum moleculare, ML - molecular layer of the dentate gyrus, GCL - granular cell layer of the dentate gyrus, PL - polymorphic layer (hilus) of the dentate gyrus*. B) Western blot images and depicting the average expression of glycosylated (50 kDa) and aglycosylated (37 kDa) Kir4.1 proteins in the hippocampus (left) and somatosensory cortex (right) of young and aged 5xFAD mice. C-F) Box plots depicting the relative expression of glycosylated (50 kDa; C, D) and aglycosylated (37 kDa; E, F) Kir4.1 proteins in the hippocampus (C, E) and somatosensory cortex (D, F).

### The impact of astrocytic K^+^ clearance on firing properties of nearby neurons

The extracellular K^+^ concentration is one of the key players determining neuronal signalling, including the passive and active properties of neurons (7, 8). We therefore assessed the functional impact of local alterations in astrocytic K^+^ clearance rate at the SLM layer of the hippocampus on the physiological properties of nearby neurons (Fig. 6). Analysis of the resting membrane potential indicated a more depolarized resting membrane potential in 5xFAD mice, which was due to both ageing and 5xFAD mutations: *F*(*_1,42_*) = 9.81, *p* = 0.003 for the factor ageing; *F*(*_1,42_*) = 8.52, *p* = 0.005 for the factor 5xFAD mutations; *F*(*_1,42_*) = 0.15, *p* = 0.69 for the factor interaction, two-way ANOVA, Fig. 6A). While both input resistance and time constant were comparable between groups (Fig. 6B, C), the spike rheobase (minimal current required to elicit an action potential) was significantly affected by both the ageing process and 5xFAD mutations: *F*(*_1,41_*) = 11.31, *p* = 0.001 for the factor ageing; *F*(*_1,41_*) = 10.43, *p* = 0.002 for the factor 5xFAD mutations; and *F*(*1,41*) = 0.21, *p* = 0.64 for the factor interaction; two-way ANOVA. On average, the rheobase level of neurons in the aged 5xFAD mice was significantly decreased compared to aged WT mice (90.35 ± 6.3 pA, *n* = 14 vs 125.45 ± 13.75 pA, *n* = 11; respectively, *p* < 0.01, two-way ANOVA with Sidak’s post hoc test, Fig. 6D). Moreover, there was a significant increase in the rheobase level in the aged WT group vs the young WT group (125.45 ± 13.75 pA, *n* = 11 vs. 89.09 ± 8.41 pA, *n* = 11, respectively, *p* <0.05, two-way ANOVA with Sidak’s post hoc test), indicating the impact of normal ageing on the excitability profile of these neurons. This data supports the previously revealed trend that while physiological ageing decreases the excitability of neurons residing in the SLM layer of the hippocampus, 5xFAD mutations result in the development of a high level of neuronal excitability (53, 54).

**Figure 6.**
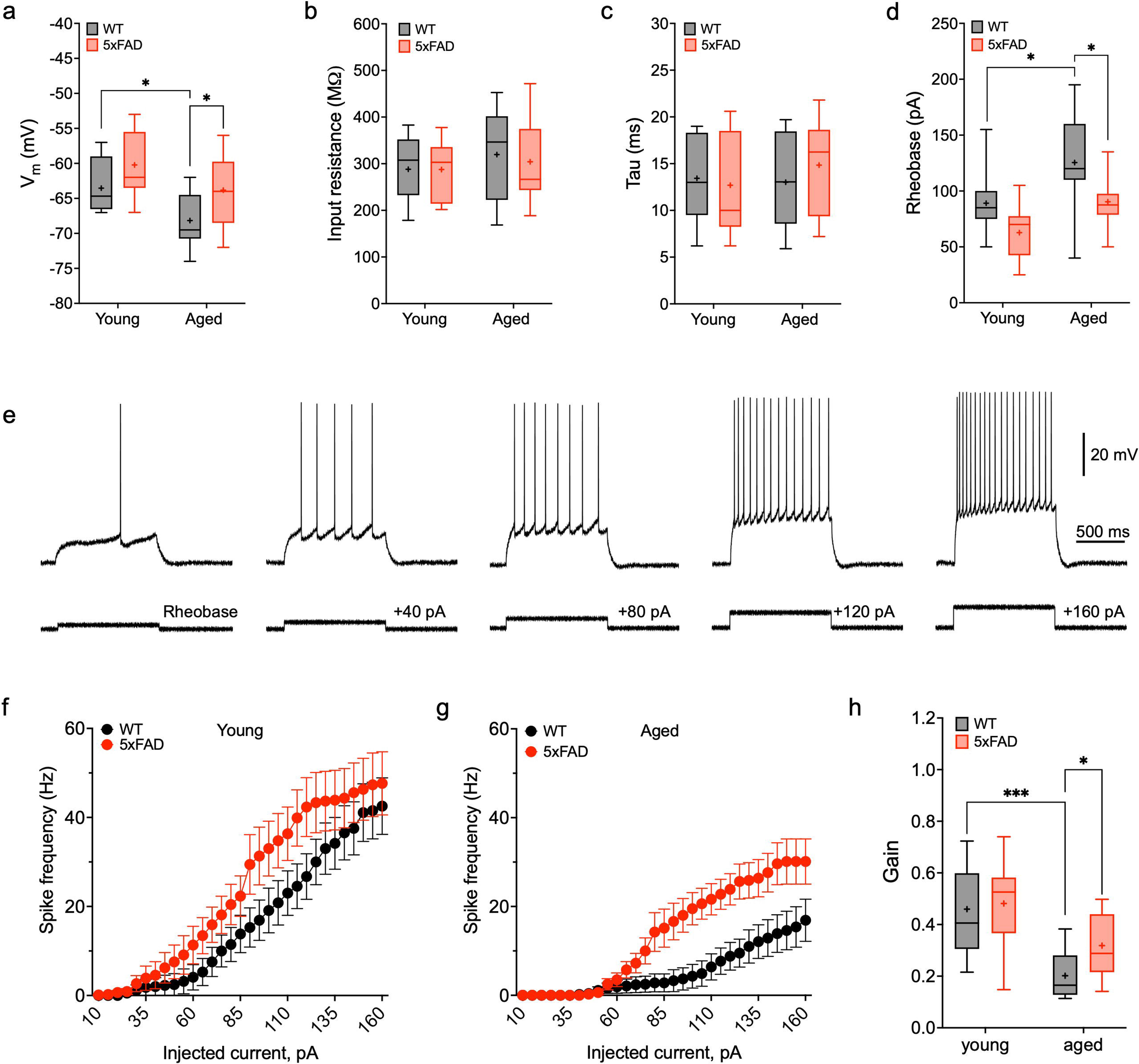
Membrane properties of neurons recorded from the stratum lacunosum moleculare layer of CA1 area. A-D) Box plots depicting the average RMP (A), Input resistance (B), Time constant (C) and spike rheobase (D). E) Sample traces of spiking neurons during injections of increasing step currents. F, G) F-I plots depicting the correlation between the increase in injected stimulus current and the spiking frequency in young (F) and aged (G) mice. H) Box plot depicting the overall gain of the F-I curves in both 5xFAD and WT mice.

Repetitive firing is a fundamental feature of spiking neurons, which determine their overall response to synaptic inputs during activation (55). We next assessed neuronal firing properties following injections of increasing step currents (5 pA steps for 1 s, Fig. 6E). Analysis of the F-I curves indicated a significant increase in the gain of the F-I curves, which was mainly due to aging and to less extent the 5xFAD mutations (*F*(*_1,33_*) = 22.06, *p* < 0.0001 for the factor ageing; *F*(*_1,33_*) = 2.85, *p* = 0.1 for the factor 5xFAD mutations; *F*(*_1,33_*) = 1.49, *p* = 0.23 for the factor interaction, two-way ANOVA, Fig. 6F-H). Indeed, posthoc analysis indicated a significant decrease in aged mice from both WT (0.182 ± 0.02, *n* = 9 in aged WT vs 0.46 ± 0.04 in young WT, *n* =11, *p* > 0.001), and 5xFAD mice (0.482 ± 0.06, *n* = 9 in young 5xFAD vs 0.318 ± 0.4 in aged 5xFAD, *n* = 8, *p >* 0.05, two-way ANOVA with Sidak’s post hoc test). Overall, these data indicate a decline in neuronal excitability with ageing, as evidenced by a reduction in the slope of F-I curves. Meanwhile, the presence of 5xFAD mutations appears to maintain a high cellular excitability profile.

K^+^ conductance plays a key role in shaping neuronal signals, specifically as it increases at the repolarisation phase of the action potential (56). Hence, alterations in [K^+^]_o_ clearance rate may result in significant changes in spike shape, specifically the half-width spike amplitude (HWSA) (7). To assess the impact of altered [K^+^]_o_ clearance rate in the hippocampus of aged 5xFAD mice, we have compared the HWSA of neurons during a burst of action potentials which significantly increase [K^+^]_o._

To reduce the effect of spike shape variability between neurons, we normalised the HWSA of the 5^th^ spike to the HWSA of the spike rheobase of each neuron (Fig. 7A). Our data indicate that in all neurons recorded, an increase in the firing frequency of neurons led to a significant rise in the HWSA of all neurons (Fig. 7B, C). However, there was a significant effect of 5xFAD mutations on the rise of HWSA: *F* (*_1,32_*) = 2.78, *p* = 0.1 for the factor *ageing*; *F*(*_1,32_*) = 4.72, *p* > 0.05 for the factor *5xFAD mutations*; *F*(*_1,32_*) = 0.64, *p* = 0.42 for the factor *interaction*). On average, the gain in HWSA was significantly higher in the aged 5xFAD mice vs the aged WT mice (0.54 ± 0.05, *n* = 12 vs 0.3 ± 0.03, *n* =7, respectively, *p* > 0.05, two-way ANOVA with Sidak’s post hoc test, Fig. 7D). Also, the HWSA of neurons from young WT group was higher than in the aged WT mice (0.5 ± 0.07, *n* = 10 vs 0.3 ± 0.03 *n* = 7, respectively, *p* = 0.09, two-way ANOVA with Sidak’s post hoc test). These findings indicate an increased accumulation of K^+^ in the extracellular space following neuronal firing in the aged 5xFAD mice. Additionally, the decrease in HWSA found in the aged WT mice implies an increase in the [K^+^]_o_ clearance rate during the maturation of the glial cell syncytium, which is consistent with recent in vivo measurements (57).

**Figure 7.**
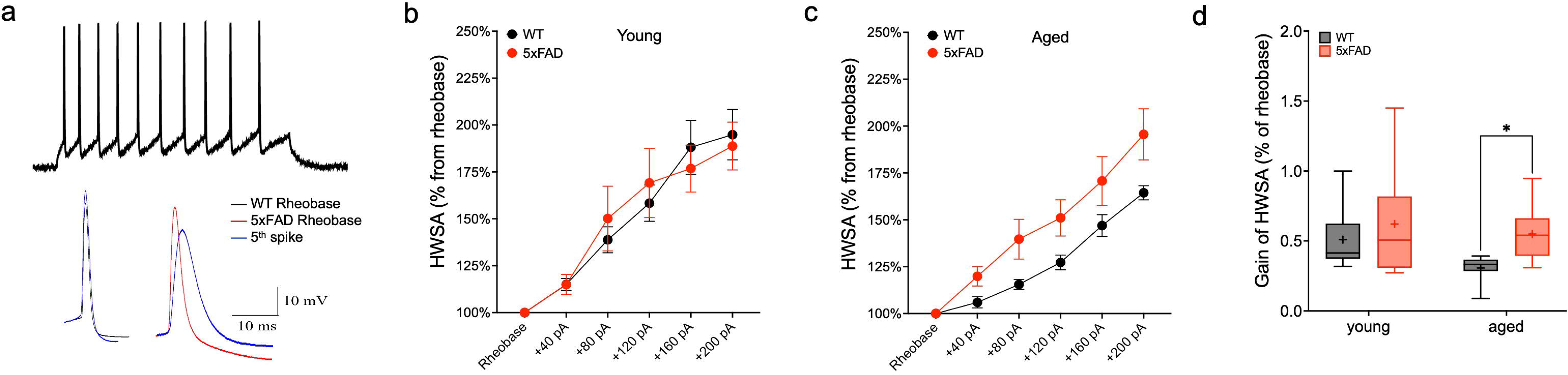
Alterations in Astrocytic K^+^ clearance affect neuronal signalling. A) sample traces of a spike burst (top) and individual spikes (bottom) depicting the shapes of the spike rheobase and the 5^th^ action potential in a spike train (stimulus amplitude of 200 pA) in aged WT (left) and aged 5xFAD (right) mice. B) Dot plot depicting the average HWSA recorded from young (B) and aged (C) mice of both 5xFAD and WT littermates. D) Box plot depicting the average increase in the 5^th^ HWSA compared to the HWSA of the 1^st^ spike in the spike train.

## Discussion

New insights into the genetic etiology of AD highlight the role of glia in the pathophysiological process (58). Indeed, transcriptomic analysis of astrocytes in AD patients and mouse models revealed the upregulation of genes involved in immune signalling pathways and the downregulation of genes related to neuronal support function, indicating a potential impairment of astrocytes in providing optimal support and maintenance to neurons in AD (59, 60).

Previous studies indicated on Na^+^ and K^+^ ion imbalances in AD (61), and a very recent paper indicated that in a mouse model for AD the baseline extracellular concentration of K^+^ is higher than in age-matched control (57). In this study, we measured both extracellular and intracellular K^+^ dynamics during disease progression in a mouse model for AD. Our data indicate a significant decrease in the astrocytic [K^+^]_o_ clearance rate in the hippocampus of aged 5xFAD mice, which was region-specific, as no significant alterations were detected in the superficial layers of the somatosensory cortex.

These impairments in [K^+^]_o_ clearance rate were mediated by a decrease in K^+^ conductance via Kir_4.1_ channels, which play a major role in K^+^ uptake (Fig. 3). Indeed, Kir channels are widely distributed in glial cells (50), and knockdown of these channels using RNA interference has been shown to disrupt transmembrane K^+^ flow induced by extracellular KCl application (62). Hibino and colleagues show that there is region-specific distribution of Kir4.1 and Kir5.1 channels, as the heteromeric Kir4.1/5.1 assembly was widely distributed in the cortex while the homomeric Kir4.1 was mainly distributed in the hippocampus and thalamus (63). Hence, the impact of these channels on the [K^+^]_o_ clearance varies between different brain regions, which is consistent with our results indicating significant alterations in the hippocampus, but not the somatosensory cortex of aged 5xFAD mice (Fig. 3, 4). Interestingly, in our study of 5xFAD mice, we found a decrease of only one of the Kir4.1 isoforms, with a molecular weight corresponding to the glycosylated form of the Kir4.1 protein that is mainly expressed in astrocytes (51, 52, 63).

Our data also indicate that the alterations seen in astrocytic function were accompanied by morphological changes (Fig. 2), as astrocytes in the hippocampus of aged 5xFAD mice had a higher soma volume and diameter, as well as longer and thicker processes, both of which are characteristics of reactive astrocytes (38) and indicators of ongoing neuroinflammation. These results are in line with previous studies suggesting that misfolded proteins, such as Aβ and tau, trigger persistent neuroinflammation that affects astrocytic reactivity (11, 64, 65) and further neurodegeneration in the AD brain (25, 66, 67). These morphological changes also translated to alterations in the panglial syncytium, as astrocytes in the hippocampus of aged 5xFAD mice expressed a lower coupling ratio (Fig. 2), consistent with their impairment to clear K^+^ via spatial buffering (8, 34). Our study found different patterns of astrocytic reactivity in the hippocampus and somatosensory cortex, indicating either different sensitivity to misfolded proteins or different efficacy of the compensatory mechanisms at these brain regions. While consistent with previous studies (68), the mechanism leading to this variability is still unclear; however, the fact that astrocytic reactivity was much less evident in the somatosensory cortex point to possible differences in astrocytic heterogeneity between these regions.

Indeed, it has been shown by several research groups that protoplasmic astrocytes across all brain regions are heterogeneous (69, 70). Astrocytes in various brain regions exhibit differences at both functional and anatomical levels (71), which in turn affects their responses to different stimuli. This disparity in astrocytic composition may also impact their connectivity, as we revealed significant differences in the number of primary connected astrocytes in the hippocampus and SSC of control mice. Interestingly, a recent study comparing biochemical changes in astrocytes from the cortex and hippocampus identified significant alteration of biochemical processes in astrocytes from the hippocampus but not in the cortex of 5xFAD mice (72).

In the hippocampus, the stratum lacunosum moleculare (SLM) layer serves as a connection hub between the entorhinal cortex and CA1 region and thus crucial for memory processing. The SLM contains the highest density of GFAP-expressing astrocytes (73) as well as a large population of GABAergic interneurons and plays a critical role in modulating dynamic activity in hippocampal networks (74). Genetic deletion of Cx-43 and Cx-30 in astrocytes results in impaired spatial buffering (75) and a recent study indicated axonal loss and reduced cholinergic input from the medial septum into the SLM of a mice model for AD (76). Our data indicate that the reduced [K^+^]_o_ clearance rate in aged 5xFAD mice led to functional implications on neighbouring neurons, expressed as an increased excitability profile (Fig. 6D, G-H) from one end, and an increase in HWSA in a spike train, compatible with an increase in [K^+^]_o_. This is consistent with previous studies (61), including a very recent paper indicating that while the baseline extracellular concentration of K^+^ decreases with age, it is higher in AD mice than in age-matched control (57).

In summary, our data are consistent with previous studies indicating AD is associated with neuronal hyperexcitability (64, 72, 77, 78), and further suggest that alterations in astrocytic [K^+^]_o_ clearance rate result in abnormal neuronal network excitability as previously shown (7, 8), which can lead to hypersynchrony, aberrant oscillatory rhythmic activity, interneuron dysfunction and neuronal loss that contribute to the cognitive decline observed in Alzheimer’s disease (3).

## Data availability

The data that support the findings of this study are available from the corresponding author upon reasonable request.

## Acknowledgements

We thank Dr. Sindy Kueh for her technical assistance. This study was supported by WSU Internship to Armaan Mangat and the Ainsworth Medical Research Innovation Fund awarded to Yossi Buskila and John W. Morley. Evgeniia Samokhina was supported by the Ainsworth MRIF Scholarship Program.

